# NEuRT: A Transformer-Based Model for Explainable Neuronal Activity Analysis

**DOI:** 10.64898/2026.01.13.699305

**Authors:** Georgii Raev, Daniil Baev, Evgenii Gerasimov, Viacheslav Chukanov, Ekaterina Pchitskaya

**Affiliations:** Laboratory of Biomedical Imaging and Data Analysis, Peter the Great St. Petersburg Polytechnic University, Khlopina St. 11, St. Petersburg 194021, Russia; Laboratory of Molecular Neurodegeneration, Peter the Great St. Petersburg Polytechnic University, Khlopina St. 11, St. Petersburg 194021, Russia; Laboratory of Molecular Neurobiology, Pavlov Institute of Physiology, Russian Academy of Sciences, St. Petersburg 199034, Russia

**Keywords:** neuronal activity, Alzheimer disease, transformer, miniscope, MICrONS, neural networks, machine learning

## Abstract

The study of neuronal activity is essential for understanding brain function and its alterations in neurodegenerative diseases. Advances in in vivo imaging have enabled real-time observation of neuronal dynamics, but classical statistical methods struggle to capture the complex, time-dependent interactions within neuronal networks. Machine learning offers promising solutions for analyzing high-dimensional neuronal data, yet their application in neuroscience remains limited. Here, we introduce NEuRT, a Bidirectional Encoder Representations from Transformers (BERT)-based model adapted for neuronal activity analysis. NEuRT leverages self-attention mechanisms to interpret complex neuronal interactions, providing insights into patterns that traditional methods may overlook. Pre-trained on the recently introduced large annotated dataset MICrONS for signal reconstruction, NeuRT demonstrates strong generalization, effectively reconstructing activity from both visual cortex two-photon and hippocampal miniature fluorescence microscopy. Built on the BERT architecture, the NEuRT model can be efficiently fine-tuned for a wide range of downstream tasks. We showcase its application in classifying wild-type and transgenic Alzheimer’s disease model mice, based on hippocampal activity, revealing group-specific features through attention map analysis. By reducing reliance on extensive labeled data, addressing a critical challenge in neuroscience, NEuRT bridges fundamental neuroscience and disease research, offering a robust framework for AI-driven and explainable neuronal activity analysis.

## I. Introduction

THE study of neuronal activity is a cornerstone of modern neuroscience, providing critical insights into the neuronal function and its relationship to behavior. Advances in *in vivo* imaging – particularly through two-photon microscopy and the more recent development of miniature fluorescence microscopy, have revolutionized the researcher’s knowledge in real-time brain neuronal dynamics. Miniature fluorescence microscopy, in particular, enables imaging in freely moving animals, offering unprecedented opportunities to investigate neuronal circuits under naturalistic conditions [1], [2]. By analyzing activity patterns across brain regions, researchers could uncover fundamental principles of neuronal function and identify disturbances in neuronal networks associated with neurodegenerative diseases, such as Alzheimer’s disease. These tools not only deepen our understanding of the fundamental principles of brain functioning at the network level, but also pave the way for new insights into the pathogenesis and development of neurological therapies.

Classical statistical methods are commonly used to analyze neuronal activity data [3]. Dimensionality reduction techniques are employed to examine the relationship between behavior and brain activity [4], [5]. Moreover, variuos approaches enable registering cells across multiple sessions [6], with subsequent identification and analysis of neuronal ensembles [7], [8]. However, classical statistical approaches often fail to account for complex and time-dependent interactions between neurons, which potentially could be investigated with machine learning methods.

Machine learning-based solutions have proven themselves well for analyzing structurally complex and multidimensional data, which is rightfully for neuronal activity datasets. Machine learning techniques have been applied [9] for classification of mouse behavioral states based on the mesoscopic cortex-wide calcium imaging data. Moreover, there is growing interest in leveraging large foundation models for various neurobiology applications [10]. However, the effectiveness of these approaches-particularly foundation models-hinges on the availability of large-scale, well-annotated datasets, which remain scarce in animal-based biological research. Initiatives such as the MICrONS project, which combines two-photon microscopy of the mouse visual cortex with detailed visual stimulus annotations, have provided unprecedented insights into neuronal circuit organization [11], and gave the researchers access to the biggest available to the date dataset.

However, analyzing such multidimensional and large-scale data continues to pose significant challenges. For example, using the MICrONS dataset, a foundation model was trained to predict cortical neuronal activity in response to visual stimuli [12]. Numerous transformer-based architectures and attention mechanisms are employed to analyze medical data [13], including multimodal models for diagnosing neurodegenerative diseases [14]. Transformers are particularly promising due to their potential for interpreting results, allowing for more precise analysis of complex neuronal interactions [15]. However, transformer-based architectures have so far been used rather sparsely for tasks related to the study of neuronal activity [16].

In this paper, we introduce the NEuRT model, a Bidirectional Encoder Representations from Transformers (BERT)-based architecture adapted for neuronal activity analysis. Originally developed for natural language processing, the BERT architecture leverages self-attention mechanisms to capture complex relationships within sequential data. A key advantage of NEuRT is its interpretability: the self-attention mechanism enables the model to weigh the significance of neuronal events relative to one another, revealing patterns and correlations that may elude linear or unidirectional models. This will make it possible in the future to identify various patterns in the work of neurons, including those inherent in neurodegenerative diseases. The NEuRT was pre-trained on visual cortex two-photon microscopy data from the MICrONS dataset for signal reconstruction tasks. Notably, the model generalizes effectively, successfully reconstructing activity from alternative sources and brain areas, particularly dorsal hippocampal neuronal activity recorded via miniature fluorescence microscopy method. BERT models can be pretrained on large datasets of neuronal activity and be fine-tuned for specific tasks. This reduces the need for large labeled datasets, which are often scarce in neuroscience. The present manuscript demonstrates the successful application of NEuRT for classification tasks, specifically in distinguishing between wild-type and transgenic Alzheimer’s disease model mice based on hippocampal neuronal activity recorded using miniature fluorescence microscopy. Subsequent interpretation of the model’s attention maps and other analytical techniques further revealed group-specific neuronal features, offering novel insights into Alzheimer’s-related neuronal dysfunction. Thus, NEuRT provides a robust framework for an explainable artificial intelligence (AI)-driven neuronal activity analysis, bridging computer science and fundamental neuroscience.

## II. Materials and methods

### A. Animals

Transgenic 5xFAD mice (Jackson Laboratory, USA; strain #034848), carrying five familial Alzheimer’s disease (AD) mutations and exhibiting intraneuronal β-amyloid accumulation, neurodegeneration, and neuronal loss [17], were used for in vivo recordings of neuronal activity. These mice, maintained on a C57BL/6 genetic background, were bred and housed at the Laboratory of Molecular Neurodegeneration, Peter the Great St. Petersburg Polytechnic University. Mice were kept in groups of 4-6 per cage under standard vivarium conditions (12-hour light/dark cycle) with ad libitum access to food and water. Experiments started at 6,5 months old when significant memory loss, β-amyloid accumulation in the brain and synaptic deficiency were observed in transgenic 5xFAD mice [18]. All experimental procedures complied with the ethical standards outlined in the European Convention for the Protection of Vertebrate Animals used for Experimental and Other Scientific Purposes (Strasbourg, 1986) and the Declaration of Helsinki (1996) on the humane treatment of animals.

### B. Viral constructs delivery

Mice were initially placed in an induction chamber and anesthetized with 2.0% isoflurane. Once the animal ceased voluntary movement, it was transferred to a stereotaxic frame (Model 68001, RWD Life Science, China), where anesthesia was maintained with continuous isoflurane delivery (1.5% - 2.0%) throughout the entire surgical procedure. Surgery began only after the animal failed to respond to a paw pinch, confirming adequate level of anesthesia. Throughout all manipulations, mice were positioned on a heated mat equipped with a temperature controller (Model 69002, RWD Life Science, China), with the temperature maintained at 37 °C. A unilateral injection of the viral vector AAV5.Syn.GCaMP6f.WPRE.SV40 [19] was performed into the left hemisphere. The injection coordinates relative to bregma were: anteroposterior (AP) – 2.1 mm, dorsoventral (DV) – 1.45 mm, and mediolateral (ML) +1.4 mm. A total volume of 1.15 µL was delivered at a rate of 0.1 µL/min using a Hamilton syringe (#84853, 7758-02, Hamilton, USA), with a final viral titer exceeding 1 × 10^13^ vg/mL. Following the injection, the syringe was left in place for an additional 10 minutes to allow diffusion of the viral construct within the dorsal hippocampus. Afterward, the syringe was carefully withdrawn, the scalp was sutured using sterile surgical thread, and the wound area was disinfected with betadine.

### C. GRIN lens implantation and baseplates fixation

Three weeks post injection, GRIN lens implantation was performed. Mice were preliminary anesthetized (2.0% - 2.5% of isoflurane) in the induction chamber and moved to the stereotaxic apparatus. The mouse’s head was secured in the stereotaxic frame, and the scalp was removed with fine surgical scissors to expose the skull. The skull surface was cleaned with 3% hydrogen peroxide. Then, a skull was carefully scratched for a future cement better adhesion. A 2-mm diameter craniotomy was drilled (Strong 90n, SAESHIN PRECISION CO, South Korea) above the injection site. Cortical tissue was carefully aspirated under continuous perfusion with phosphate-buffered saline (PBS) until vertical fibers of the corpus callosum became clearly visible. GRIN lens implantation (#64519, Edmund Optics, USA) was implanted only after bleeding had ceased and the corpus callosum remained clearly visible for at least 3 minutes. A small vent was then secured to the right lateral skull. The GRIN lens was slowly lowered to a depth of 1.45 mm from the medial edge of the craniotomy and fixed by a small drop of cyanoacrylate glue. The entire exposed skull was then sealed using light-curing dental cement (Dent-Light Flow, tdVladmiva). At the end of the surgery, mice received intraperitoneal injections of 50 µL of atipam and 1 mg/kg dexamethasone to promote recovery and reduce inflammation. Postoperative recovery after GRIN lens implantation lasted approximately 4–6 weeks.

Following recovery, mice were anesthetized with isoflurane (1.5% - 2.0%) and transferred to stereotaxic. A baseplate for imaging was implanted to achieve the best field of view (FOV) of hippocampal neurons. When the best FOV was achieved, the baseplate was fixed using light-curing dental cement. In cases where neurons were not clearly visible, the baseplate was positioned based on the clearest vascular patterns. Mice were allowed to recover for an additional 2 weeks after baseplate placement before calcium imaging sessions started.

### D. Hippocampal neuronal activity recordings under freely behaving conditions

To habituate the animals to the recording setup, the miniscope was attached to the baseplate on the mouse’s head for 5 minutes per day over two consecutive days within the experimental environment. Recordings of neuronal activity in freely moving mice were conducted in a round arena with a diameter of 63 cm. To avoid anesthesia-induced suppression of neuronal activity and GCaMP6f fluorescence, the miniscope was mounted onto the baseplate without the use of isoflurane or any other anesthetic agents. Neuronal activity was recorded over four consecutive days, with each recording session lasting 7 minutes under identical environmental conditions and without external behavioral interference. After each session, the arena was thoroughly cleaned with 70% ethanol and allowed to dry before placing the next mouse for recording.

### E. Processing of miniscope recordings

Miniscope data was recorded using the Pomidaq (Portable Miniscope Data Acquisition) software at a rate of 15 frames per second. Excitation parameters were adjusted individually for each mouse, ensuring they did not exceed 50% of maximum intensity, while the gain setting remained at “medium” for all mice in Pomidaq version 0.4.5. First and the last minute of each session were cut, so individual recording was 5-minute long. For processing the miniscope data, we used Minian, an open-source tool for miniscope data analysis [20]. Minian performs background fluctuation elimination, motion correction, and calcium signal extraction using the CNMF method. The following parameters were applied in Minian: “wnd size” set to 1000, “method” set to “rolling”, “stp size” set to 500, “max wnd” set to 15 and “diff thres” set to 3. All the CNMF parameters were kept default except “sparse penal” in “param first temporal” and “param second temporal” which was set at 0.18.

### F. Datasets

In the present study, two datasets were utilized. The first dataset was obtained from the open-access MICrONS project and consists of functional neuronal activity data from the visual cortex acquired using two-photon microscopy. The second own dataset (hereafter referred to as the “Miniscope” dataset) was generated using miniature fluorescence microscopy and contains recordings of hippocampal neuronal activity in both wild-type (WT) mice and 5xFAD transgenic mice aged 6.5 months a widely used genetic model of Alzheimer’s disease (AD).

#### 1) MICrONS dataset

The Machine Intelligence from Cortical Networks (MICrONS) project represents one of the most comprehensive open-access resources, containing a dataset the functional imaging dataset includes recordings from approximately 75000 pyramidal neurons, capturing their responses to diverse visual stimuli [11]. For the present study, we utilized neuronal fluorescence data reflecting neuronal activity from the functional imaging component of the MICrONS dataset during the model pretraining stage. A total of 142130 masks from the dataset were used, obtained in 120 scanning fields with a unique combination of *session_idx, scan_idx*, and *field_idx* fields. The dataset volume after all the preprocessing stages was ∼ 270 gb.

#### 2) “Miniscope” dataset

We additionally employed our own dataset, which contains recordings of hippocampal neuronal activity acquired using miniature fluorescence microscopy. Data were collected from freely moving mice during experiments conducted in a free moving behavior in the rounded arena. A total of 36 recordings were collected for two groups of mice: 23 sessions for 7 wild-type (WT) mice and 16 sessions for 5 transgenic Alzheimer’s disease model mice (5xFAD). Supplementary Table S1 shows detailed information about the number of neurons recorded in each session.

#### 3) Neuronal activity matrix

Neuronal activity – defined as fluorescence signals, in particular calcium traces - was supplemented with activity statistics from spatially defined neuronal groups, following the approach described in [16]. Specifically, for each neuron, the internal group was defined as neurons located within a Euclidean distance of Δ (Δ = 100 px, that equals approximately 100 µm), whereas the external group comprised neurons located at distances greater than Δ. For each group, the mean and standard deviation of neuronal activity were calculated. Consequently, the activity of each neuron was represented as follows:

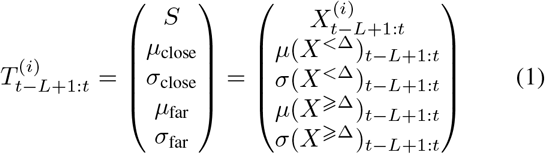

Where 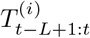 is the activity matrix of the neuron with index *i* over the time window from *t* − *L* + 1 to *t*, where *L* is the fixed number of frames. *X*^(*i*)^ is the signal of the neuron *i*, while *X*^*<*Δ^ and *X*^⩾Δ^ correspond to the signals of neurons located at distances less than and greater than Δ from the neuron under consideration, respectively. *µ* and *σ* denote the mean and standard deviation operations.

This representation addresses the variability in the number of neurons recorded across different sessions while incorporating the activity of the entire neuronal population. Furthermore, it is invariant to the ordering of neurons during processing. On the first step of the preprocessing, the data was resampled to a single sampling rate (15 fps). This step is necessary because the MICrONS dataset contains data with varying sampling rates that differ from the sampling rate of our own “Miniscope” dataset. Then, Min-max normalization of neuronal activity over the entire session was performed:

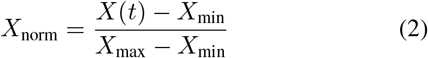

Where *X*(*t*) is the neuronal activity at time step *t, X*_min_ is the minimum activity value among all neurons in the session, *X*_max_ is the maximum activity value among all neurons in the session, and *X*_norm_ is the normalized activity value at time t. It is needed for scaling all data to the same range of values while preserving the relative magnitude of neuronal activities. For the MICrONS dataset, an additional step was performed before this normalization: negative values in the fluorescence data were replaced with zero. On the next step, for each neuron, a permutation-invariant representation was constructed, resulting in a 5-component vector for each time step (frame). Time-ordered vectors form a feature matrix. On the last step, the activity in the preprocessed form is divided into slices of *L* = 512 frames with random shifts. As a result, a 512 × 5 feature matrix is obtained, which serves as the input to the model. For each session, *n* · *k* such matrices are formed, where *n* is the number of neurons, and *k* is the number of slices for a specific dataset.

### G. NEuRT neural network architecture

#### 1) Architecture

We employed a BERT-based architecture adapted for time-series analysis, incorporating a multi-head self-attention mechanism. The model input is an activity matrix of size *L* × *S*_dim_, where *S*_dim_ denotes the number of input features (*S*_dim_), *L* is length of time-series. Instead of the token embedding layer typically used in BERT models, the data first pass through an embedding block specifically designed for time-series input, implemented as a linear transformation (one linear block with added learned positional encoding). The resulting representation is a matrix of size *L* × *h*_model_, where *h*_model_ is the hidden dimension of the model. The encoder consists of a standard BERT transformer block, modified for the changed hidden dimension of the model. In particular, the position-wise feed-forward (PFF) layer has a dimension of *h*_ff_ = 4 · *h*_model_. The number of heads in the attention layers is equal *H*. The number of encoder blocks is equal *N* . These parameters have been adapted to our task. Residual connection with layer normalization used as in original BERT. The model architecture differs slightly between the pretraining and fine-tuning stages. During pretraining, random masking is applied to the input prior to model processing, and a single linear layer is used as a decoder to reconstruct the activity matrix of size *L* × *S*_dim_. During fine-tuning, the decoder is replaced with a task-specific classifier and masking is omitted.

#### 2) Pretrain

For model pretraining, the task was formulated as masked sequence reconstruction using the MICrONS dataset. This approach is analogous to the masked language modeling objective commonly employed in natural language processing, where models learn fundamental structural patterns by predicting missing tokens [21]. In this current manuscript, this strategy enables the model to capture the intrinsic temporal and statistical properties of neuronal activity. To prevent the emergence of abnormally large weights, the model weights were initialized using the Xavier method to ensure a moderate variance balance [22]. Prior to model input, data were processed by a masking block, implemented following the SpanBERT masking strategy [23] and adapted for multidimensional time-series data. Masking was applied along the temporal axis such that all features of a given time step were masked simultaneously. The span length *n* (i.e., the number of consecutive masked time steps) was sampled from a geometric distribution biased toward shorter spans: *P* (*n*) = *p*(1 − *p*)^*n*−1^, *p* = 0.2, with a maximum n-gram length of 10. Starting indices for masking were drawn from a uniform distribution across the sequence. In total, 15% of the sequence elements were masked. Each value was replaced with zero with probability 0.8, and with a random value sampled uniformly from the interval [0, 1) with probability 0.2. This procedure encourages the model to learn to infer missing temporal segments while remaining robust to both complete dropout and noisy signal perturbations.

The loss function used was the mean squared error (MSE), applied to the masked portion of the input activity matrix:

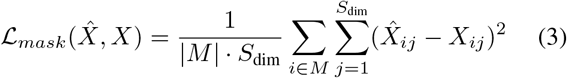

Where *M* – are indices of masked frames and | *M* | – their count. *X* is input activity matrix and 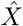 is output reconstruction of it.

As the base, we chose a model consisting of *N* = 4 encoder blocks and *H* = 4 attention heads in each attention layer. The hidden size of the model was set to *h*_model_ = 128, while the size of the feed-forward layers was *h*_ff_ = 512. We used Adam [24] as optimizer with parameters *β*_1_ = 0.9, *β*_2_ = 0.999, *ε* = 10^−8^. All layers were trained with a common learning rate of *lr* = 10^−3^, and an exponential scheduler with a decay factor *γ* = 0.9 was used for stabilization. Training was conducted for 20 epochs with a batch size of *b* = 2048. To avoid the problem of data leakage during training, the test set was taken from different scanning fields [11]. The total number of matrices for each sample |train| = 11411100, |validation| = 1267900, |test| = 1529800.

The model was fine-tuned for classification task in two cases. In the first case, we aimed to show that the model can perform reconstruction and classification from a single latent space. Therefore, a classifier is added to the decoder used during the pretraining stage. The decoder weights are not updated at this stage, but the gradient flows through it to act as a regularizer. The classifier architecture consists of a single attention layer combined with a linear layer with weights 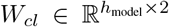. The classifier receives as input a matrix of size *L × h*_model_, which is transformed into logits according to the following formula:

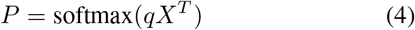

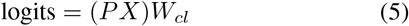

The attention mechanism is implemented using trainable parameters 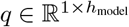. This mechanism acts as an aggregator of all hidden vectors from the encoder output, resembling a pooling operation. The pooling vector (*P* ) itself is directly involved in the model interpretation

In the first case, the model restructures the latent space to solve the second task; therefore, the loss is the sum of the cross-entropy –ℒ_*cls*_ for the classification task and regularization terms for the signal reconstruction task: ℒ*mask* – reconstruction of the masked segments, ℒ_*no*−*mask*_ – similarly, reconstruction of the unmasked segments. Thus, the final loss is represented by the following formula:

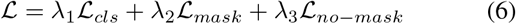

To balance the classification loss and the regularization, weighting coefficients *λ*_1_ = 1; *λ*_1_ = 1000; *λ*_1_ = 500 were used. We also doubled the maximum mask size and the percentage of masked time-steps.

The Adam optimizer is used with parameters *β*_1_ = 0.9, *β*_2_ = 0.999, *ε* = 10^−8^. Learning rate – *lr* = 10^−5^ is the same for all layers decayed to 10^−7^ with cosine annealing.

In the second case, the model was trained exclusively on the classification task, since the interpretation method described below assumes training on a single task. In this case, we removed the masking and also removed the decoder, leaving only the classifier. Cross-entropy is used as a loss function with an optimizer Adam with same parameters. In this case, the learning rate is exponentially decayed by a factor of *γ* = 0.95 at each epoch, similarly to the pretrain stage. The learning rate is set for each encoder block separately according to its depth:

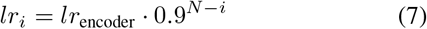

Where 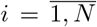, *lr*_encoder_ = 10^−6^ is the basic learning rate of the encoder. This reduction in the learning rate for deep layers is intended to improve network convergence [25]. The learning rate *lr*_embedding_ for the embedding block is equal:

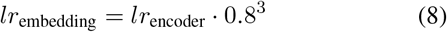

The learning rate *lr*_classifier_ = 10^−6^ is set separately for the classification block.

To avoid indirect data leakage, data was divided into training, validation and test samples based on mice in both cases. The total sample size is 88450 activity matrices: 62660 matrices for wild-type mice, 25790 for transgenic mice.

#### 3) Interpretation

To assess the importance of a specific time segment for the model’s classification decision, we employed an adapted Attention Rollout method [26]. The first step involves computing the so-called rollout from the attention maps 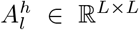, where *h* denotes the head number in encoder layer *l*:

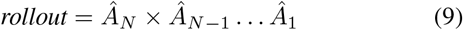

Here, *Â*_*l*_ is the head-averaged attention matrix (*A*_*l*_) from layer *l*, with the addition of the identity matrix 𝕀 to simulate residual connections:

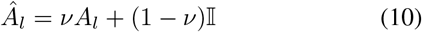

For *l* = *N*, the regular head average is used without adding the identity matrix, since it is not propagated through residual connections. The coefficient *ν* was set to 0.5, as in the original paper, reflecting the amount of information passed from one layer to the next.

In the original work, the authors used the class token column as the measure of importance, which is provided to the model in advance. In our case, such a token is absent, so we instead perform aggregation using the classifier pooling vector *P* :

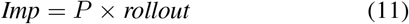

Here, *Imp* is the final vector that encodes which time segments of the input activity matrix contributed most to the decision for a particular class.

### H. Statistics

To compare distributions, the Kolmogorov-Smirnov test was applied to the cumulative sums. Statistical significance was set at *p <* 0.05. Data are presented as the mean ± standard error of the mean, unless otherwise is not specified.

## III. Results

### A. NEuRT architecture for neuronal activity analysis

In tasks related to text analysis, architectures based on the attention mechanism – Transformer, GPT (Generative Pre-trained Transformer) and BERT (Bidirectional Encoder Representations from Transformers), [27]–[29] have become widespread. The BERT architecture has demonstrated remarkable efficacy in addressing a diverse array of tasks. Among attention mechanisms, self-attention used in the BERT stands as the most well-established and thoroughly investigated approach. Additionally, architectures of this class exhibit a robust capacity to process data characterized by intricate temporal dependencies, which makes it interesting to implement it for neuronal activity analysis. The attention mechanism employed in the presented model NEuRT is independent of the direction and distance between elements in the input sequence, which is particularly valuable for neuronal activity analysis.

Encoder-only models support pre-training followed by fine-tuning for specific tasks. This approach may open new possibilities for applying AI based method in neuroscience where labeled datasets are often limited. By leveraging pre-trained representations, researchers can achieve high performance on specialized tasks while minimizing the need for extensive annotated samples, thus accelerating discoveries in brain function and dysfunction. The primary objective of this study is to develop NEuRT, a foundational model for the analysis of neuronal activity.

The overall experimental workflow is illustrated in Figure 1. Current study is based on the MICrONS publicly available dataset [11] and our own “Miniscope” dataset. Data acquired through miniature fluorescence microscopy was initially processed using the open-source minian software (Figure 1A). After data preprocessing, both the regions of interest (ROI) and neuronal activity information in the form of GCaMP6f fluorescence intensity traces (Figure 1B) were extracted. Based on these data, activity matrices consisting of five components were constructed for each neuron (Figure 1B). For the MICrONS and “Miniscope” datasets, the number of slices was set to 10 and 100, respectively, reflecting the different session durations (MICrONS: 40000-57000; “Miniscope”: 4225-4627 frames, 5 minutes). The slicing strategy was chosen to maximize session coverage while minimizing overlap between slices. The preprocessed and vectorized datasets were then used for NEuRT model training (Figure 1C). Training was conducted in two stages. First, pretraining was carried out on the MICrONS dataset using a masked sequence reconstruction task. In this stage, masked data were input into the network, which generated embedding vectors subsequently decoded back into the original sequences. The mean squared error (MSE) calculated on the masked sequence elements served as the loss function. In the second stage, a classifier producing class logits was added to the encoder. The reconstruction task served as a regularizer to prevent overfitting on the classification task. At this stage, the vectorized “Miniscope” dataset was supplemented with class labels: WT for wild-type mice and 5xFAD for the transgenic Alzheimer’s disease model mice. Model optimization was performed using cross-entropy loss with MSE used as a regularization term.

**Fig. 1.**
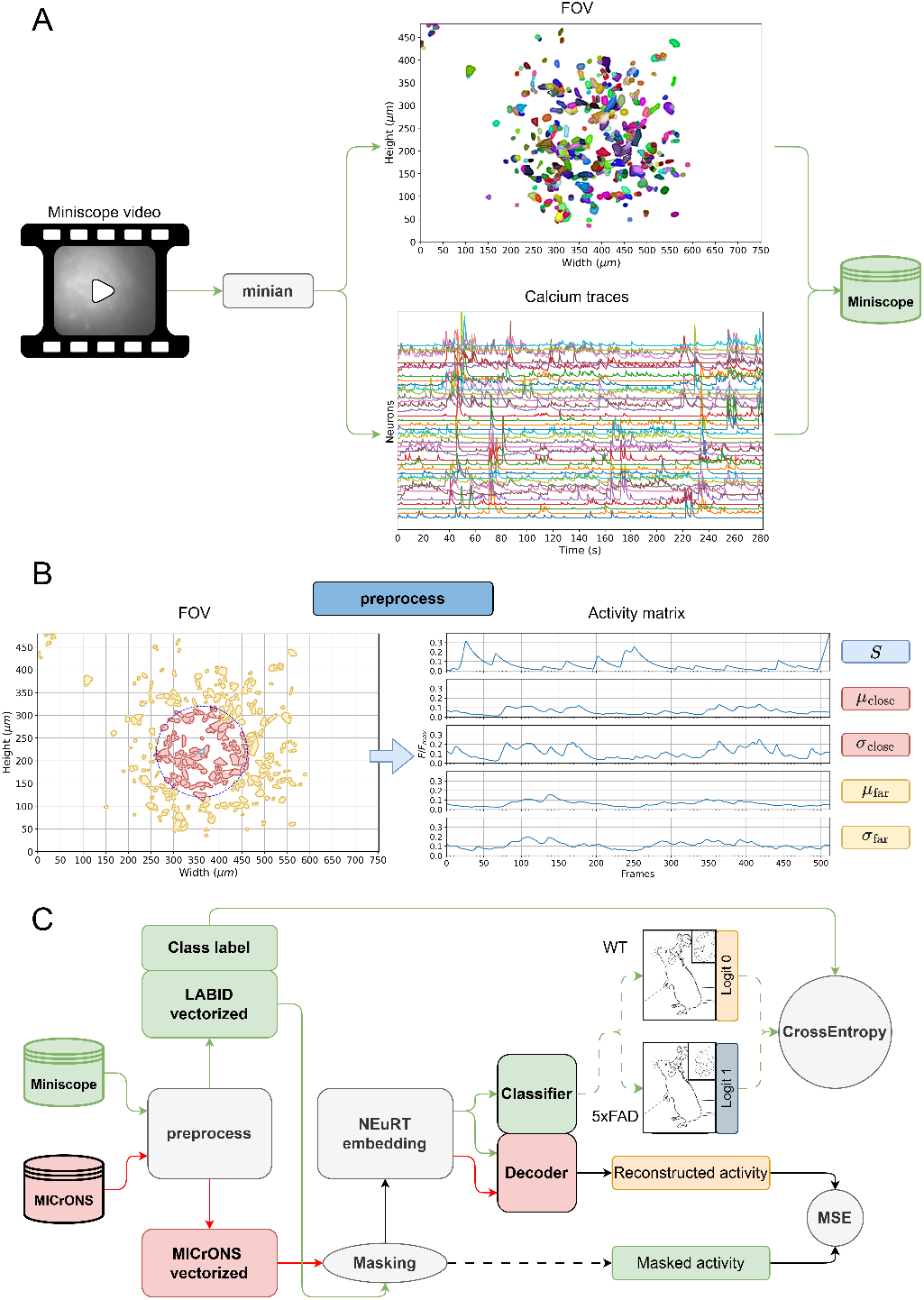
Dataflow. **A** – scheme of process miniscope video to “Miniscope” dataset. **B** – scheme of preprocessor. For each neuron separated two groups of neurons and created activity matrixes. **C** – scheme of experiment with NEuRT. Green line demonstrates dataflow for “Miniscope” dataset. The red line demonstrates data flow for MICrONS dataset. Black line demonstrates data flow for both datasets. “vectorized” is a preprocessed dataset in activity matrix format.

### B. NEuRT efficiently reconstruct the neuronal activity data from two-photon and miniscope imaging

At the first stage, the model was pretrained on the task of reconstructing masked sequences using the preprocessed MICrONS dataset. The workflow of the network during this stage is shown in Figure 2A. It consists of an embedding module, a multi-head attention encoder, and a decoder. The decoder reconstructed the activity matrix from the internal feature representation, and the mean squared error (MSE) on the masked sequence components was used as the loss function.

**Fig. 2.**
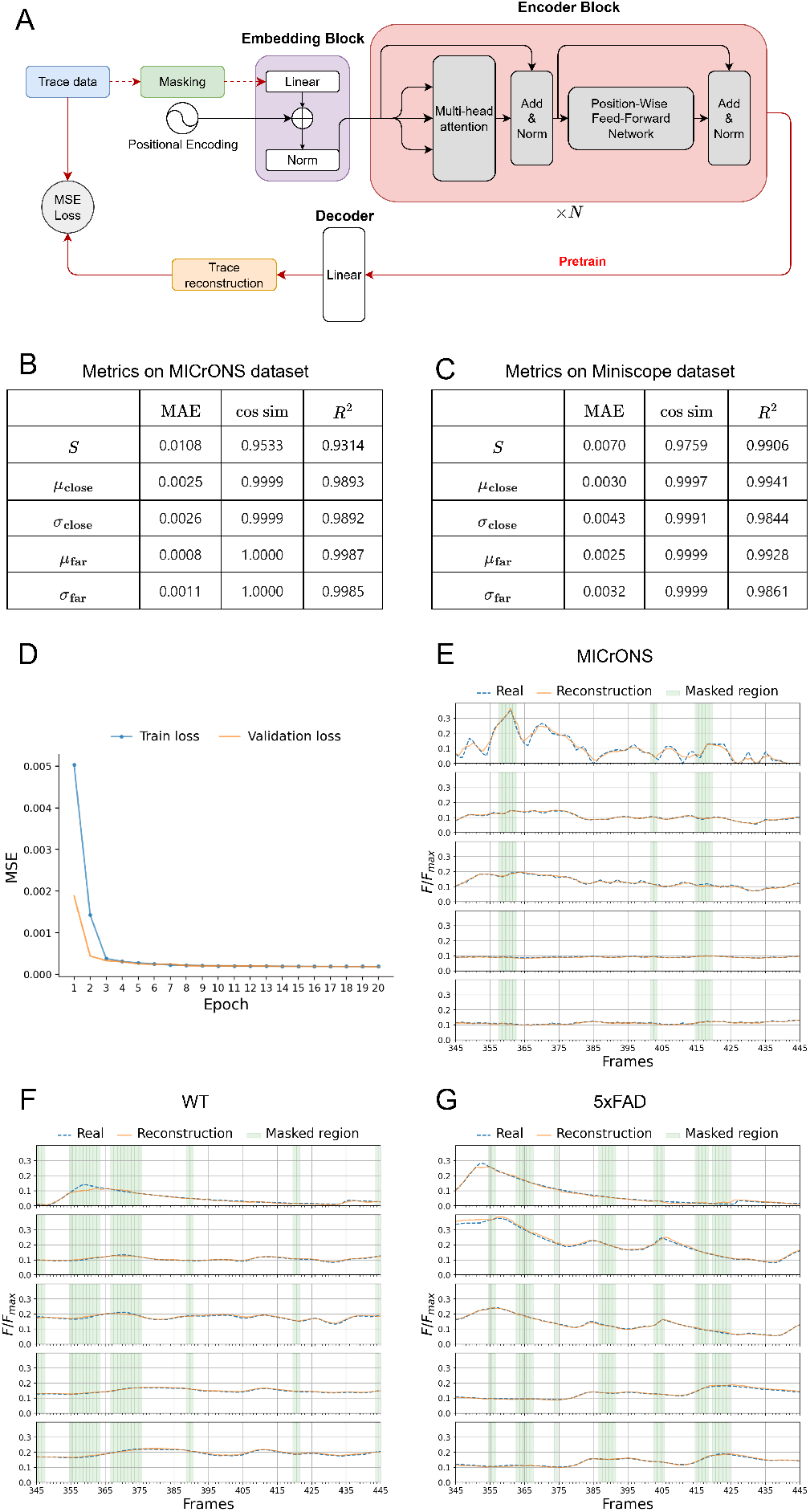
Calcium activity signal reconstruction by Neurt. **A** - scheme of NEuRT on the pretrain stage. **B** – graphic of MSE loss-function on train and validation subsets of MICrONS dataset. **C** – result of testing by activity matrix component on MICrONS test subset. **D** – result of testing by activity matrix component on “Miniscope” dataset. **E** – example of reconstructing on MICrONS test subset. **F** - example of reconstructing on wild type mouse from “Miniscope” dataset. **G** – example of reconstructing on a 5xFAD mouse from “Miniscope” dataset.

Figure 2D demonstrates the training and validation loss curves across epochs. For the training set, the plotted values represent epoch averages, which shows the initially higher loss compared with the validation set. Training was performed for 20 epochs, after which the loss reached a plateau, indicating the absence of overfitting. To further stabilize learning, an exponentially decaying learning rate was applied.

The test performance on the MICrONS dataset is shown in Figure 2B, where mean absolute error (MAE), cosine similarity, and *R*^2^ metrics are reported for each component of the activity matrix. The results indicate that the component directly reflecting neuronal activity was reconstructed less accurately (*R*^2^ = 0.9314) than the auxiliary components describing the activity of surrounding neurons (*R*^2^ = 0.9987). This reduced accuracy can be attributed to the greater variability in neuronal activity within the masked regions and the model’s tendency to smooth such fluctuations, as illustrated in Figure 2E.

Generalization ability was further evaluated on our own “Miniscope” dataset without additional training (Figure 2C). The reconstruction quality in this case remained high (*R*^2^ *>* 0.98), demonstrating that the pretrained network effectively generalized across datasets acquired with different microscopy methods. Nevertheless, reconstruction of the neuronal signal again showed lower accuracy in terms of MAE and cosine similarity compared with the auxiliary components of the activity matrix.

Examples of reconstructed signals across different datasets are presented in Figure 2E-G. Each visualization corresponds to a slice (frames 345-445) from a representative sample. Figure 2E illustrates an example from the MICrONS dataset, Figure 2F – from the wild-type (WT) mouse subset of the “Miniscope” dataset, and Figure 2G – from the mouse model of Alzheimer’s disease (5xFAD) subset of the same dataset. The plots display the ground-truth and model-predicted values for each component of the activity matrix (*S, µ*_close_, *σ*_close_, *µ*_far_, *σ*_far_), with masked regions highlighted. Legends accompanying the plots indicate all relevant designations.

### C. Joint reconstruction and classification of neuronal activity using NEuRT

In this section, we consider a downstream task for the pretrained model: binary classification aimed at distinguishing wild-type mice from 5xFAD mice – a transgenic model of Alzheimer’s disease. At this stage, a classifier head was added to the decoder block, and the loss function was defined as cross-entropy with an additional regularization term on the reconstruction task, limiting the model’s tendency to overfit to classification.

Unlike the pretraining stage, the masking strategy was modified, specifically: the maximum mask size and the number of masked time steps were both increased twofold. Cosine annealing was used to stabilize optimization. The overall architecture for this stage is shown in Figure 3A. Training was stopped when the validation classification loss reached a plateau (Figure 3B). During training, the F1 score reached a plateau on both test and validation sets (Figure 3C). To prevent data leakage, data splitting was performed at the level of individual mice. In addition, prediction accuracy was evaluated for each session within the validation set, since class imbalance limited the interpretability of accuracy computed over the entire dataset. These results are shown in Figure 3D. All the plots shown in Figure 3 reflect one of the six seeds used. Final performance showed that prediction mean accuracy with its error for 6 seeds for AD mice exceeded 98%, while the prediction error for wild-type mice was only 0.007% (Figure 3E). A comparison between the proposed model and NEuRT without pretraining, as well as baselines including logistic regression and an RNN jointly trained on classification and reconstruction tasks, is provided in Supplement Table S4. The results indicate that neuronal activity classes are linearly separable. In this case, the shortcut corresponds to the difference in mean values of the aggregated components of the activity matrices. This conclusion was drawn based on the failure of logistic regression under z-score normalization each component separately. However, both the reconstruction baseline and NEuRT without pretraining perform significantly worse on both tasks. These findings will be discussed further in the Discussion section. Cross-validation was performed to assess model stability across both tasks.

**Fig. 3.**
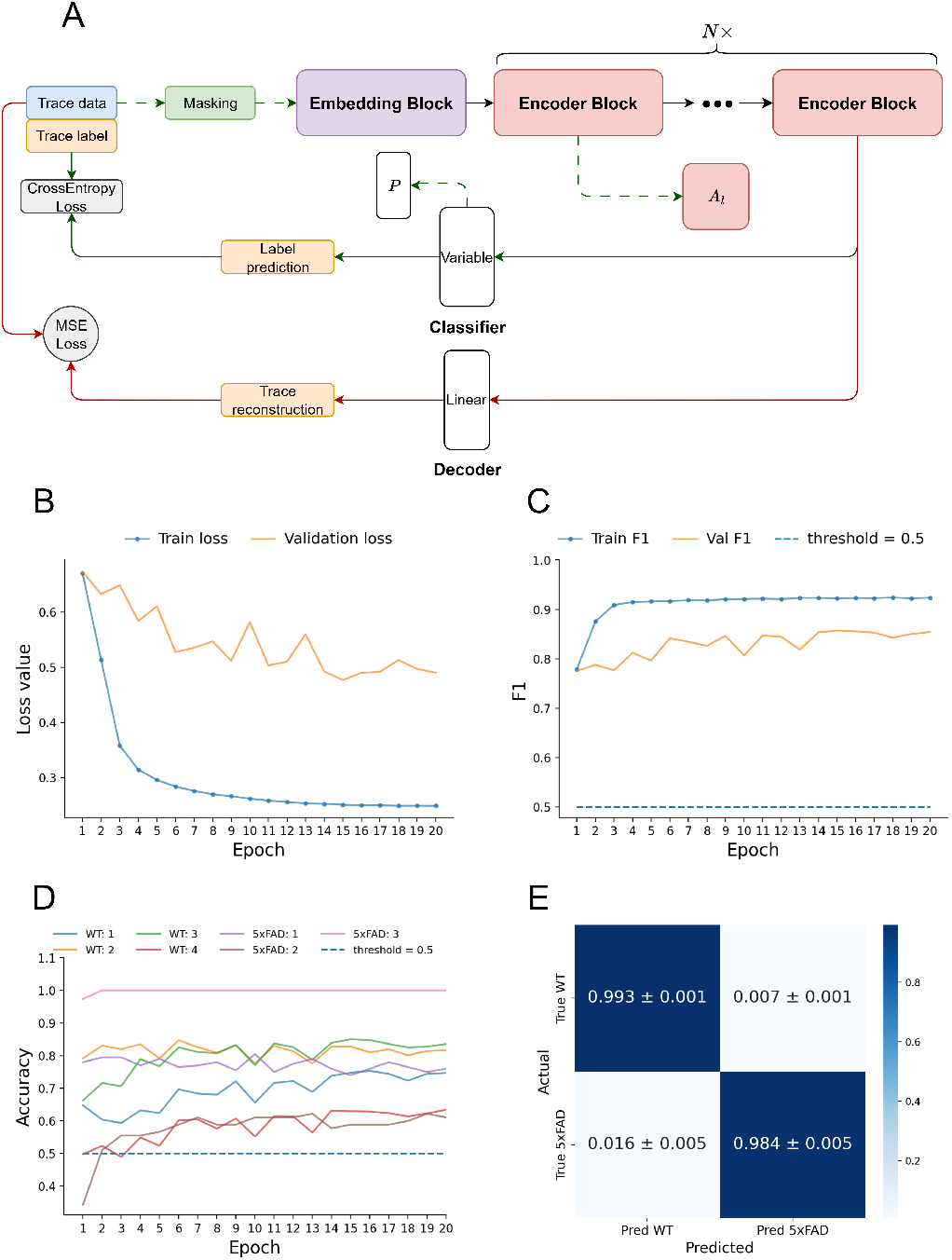
NEuRT performance in norm/pathology classification task. **A** – scheme of NEuRT on the classification stage with the demonstration of data extraction required for interpretation. **B** – graphic of cross-entropy loss-function evaluation per epoch during training on train and validation subsets of “Miniscope” dataset. **C** – graphic of F1 metric evaluation per epoch during training on train and validation subsets of “Miniscope” dataset with model threshold. **D** – accuracy evaluation per epoch during training on validation subsets of “Miniscope” dataset by each session for wild-type (WT) and Alzheimer’s disease modeling mice (5xFAD). **E** – confusion matrix on test subset on slice level with 6 different seeds of “Miniscope” dataset.

The cross-validation scheme followed that proposed in [9], where the data were split into subsets at the level of individual mice (i.e., all data from a single animal were assigned to the same subset). We define a fold as a specific partitioning of the data into training subsets. In this work, the primary focus was on training stability, the dynamics of session-level classification, and reconstruction of the activity matrix; therefore, a separate test set was not used.

The validation set consisted of data from two mice: one wild-type and one representing the Alzheimer’s disease model. Thus, each fold was uniquely defined by the pair of mice included in the validation set. During cross-validation, the mean classification performance and standard deviation were 0.8540 ± 0.2290 for the WT class, and 0.7317 ± 0.3181 for the 5xFAD class. Detailed results for each session are provided in Supplementary Table S2. The evolution of predictions for each individual session is shown in Supplementary Figure S3. In Supplementary Table S3, classification and reconstruction performance for each fold are reported. The validation set was used exclusively to monitor training dynamics and was not used to adjust the training process. This approach also reduces the total number of folds, simplifying the analysis.

### D. Interpretation of NEuRT’s attention mechanisms demonstrated neuronal activity patterns disruption associated with Alzheimer’s disease

An important objective of this work is the interpretation of the constructed model using established methods. Specifically, two issues were addressed: disentangling the influence of each component of the input activity matrix and assessing the contribution (hereafter referred to as importance) of each time segment (Figure 4). To conduct this experiment, in contrast to the fine-tuning stage, we modified the model architecture for solving only the classification task, as described in fine-tune methods. This was done to study exclusively the features used for the classification task, enabling the identification of features that distinguish wild-type from Alzheimer’s disease conditions. Importantly, while only the classifier was trained, the model became unstable to timesteps shuffling, indicating that shortcuts [30] are not the primary source of information. First, the contribution of each component of the input activity matrix was assessed. To this end, different groups of components were completely zeroed out, and the resulting matrices served as an input to the model. The predicted classes were then evaluated, and confusion matrices were constructed for each group of components. This experiment was conducted on six mice (2 wild-type and 2 Alzheimer’s disease model mice from the training set, and 1 wild-type and 1 Alzheimer’s disease model mouse from the test set), totaling over 57,000 examples. The results are presented in Figure 4B-D. They show the confusion matrices for zeroing out the mean signal values of near and far neurons, as well as the standard deviations for near and far neurons. These component groups were selected based on the changes observed in the model’s predictions when individual components were removed, as demonstrated in Supplementary Figure S4. The findings indicate that zeroing the mean values causes the model to predict almost exclusively wild-type, whereas zeroing the standard deviations leads the model to predict predominantly Alzheimer’s disease, with only 0.41% of wild-type samples classified correctly.

**Fig. 4.**
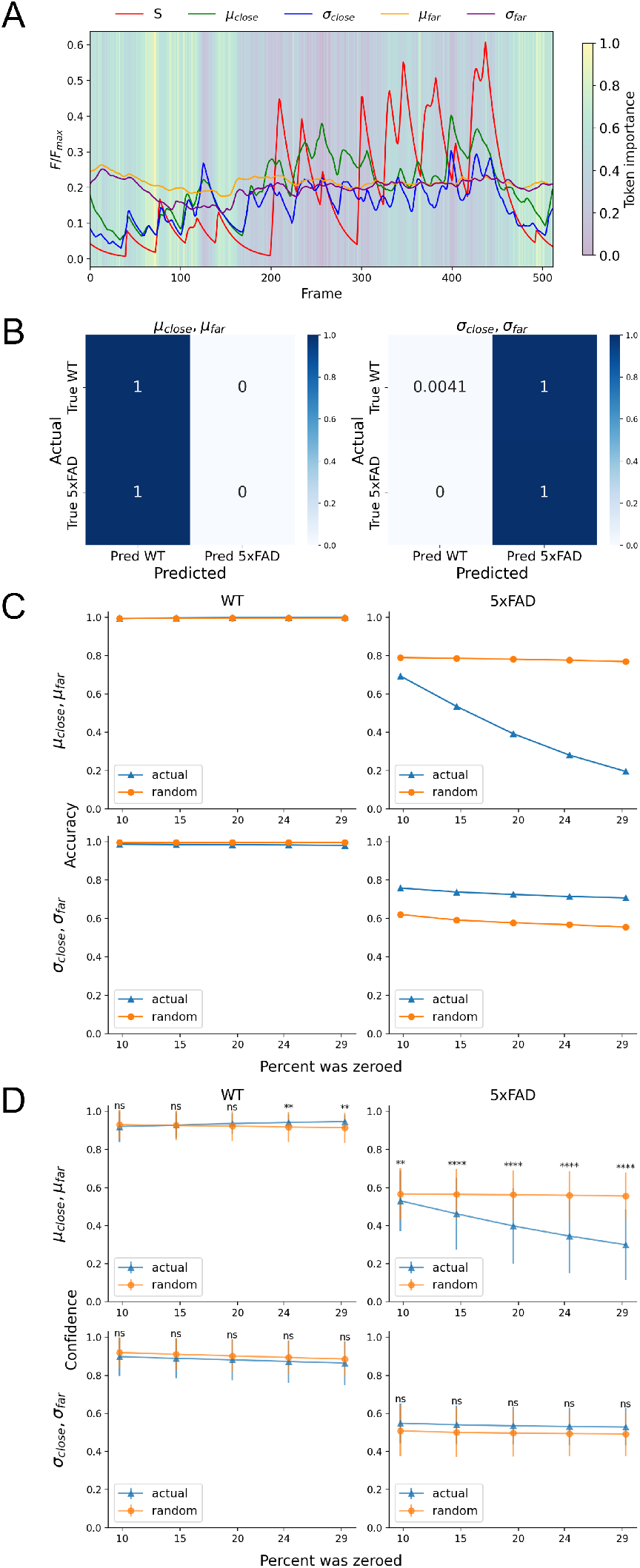
Interpretation of calcium activity signal features importance using NEuRT. **A** – visualization of the importance of each time segment, where the plot background represents importance with values ranging from 0 to 1, colored as shown on the color bar, and the curves correspond to the activity matrix values. **B** – confusion matrices for the model outputs on slice level when groups of activity matrix components are zeroed out. The color gradient on the color bar indicates results normalized to the true value. **C** – comparison of the model’s prediction accuracy for both classes when zeroing out the top-k time segments of activity matrix component groups based on their importance versus random zeroing. **D** – comparison of the model’s prediction confidence for both classes under the same conditions. Statistical comparisons were performed using the Kolmogorov-Smirnov test applied to cumulative frequency distributions: ns – non-significant; * – ***p <* 0.05**; ** – ***p <* 0.01**, *** – ***p <* 0.001**; **** – ***p <* 0.0001**

To more precisely identify Alzheimer’s disease–related patterns in hippocampal neuronal ensemble activity, we utilized the importance vector corresponding to each time segment of the input activity matrix. An example of a combined visualization of time-segment importance and the input activity matrix is presented in Figure 4A for a sample from the test set of a mouse with the Alzheimer’s disease model. In this visualization, all components of the activity matrix are displayed as curves, while the background of the plot represents the importance of each time segment, with 1 on the color scale indicating the highest importance for prediction and 0 indicating the lowest.

Based on the results described in the previous paragraph, the variability of the model prediction was evaluated when zeroing out the top-k time segments in the importance matrix within the same groups of components (mean signal values and standard deviations), according to their importance. In Figure 4C,D, the degradation of model prediction quality and the probability of assigning an example to a specific class are shown for each class separately. During the experiment, groups of components were zeroed out at time positions corresponding to the highest values in the importance vector. Specifically, by selecting a group of components, they were zeroed out at the top-10 ( ∼ 50*/*512 × 100%) to top-30 ( ∼ 150*/*512 × 100%) percent of time segments by importance, as reflected on the OX axis of Figure 4C,D, while the OY axis shows accuracy or probability of class assignment, respectively.

Based on the results above, we evaluated the variability of the model’s predictions when zeroing out the top-k time segments in the importance matrix for the same component groups (mean signal values and standard deviations), according to their importance. Figure 4C,D show the resulting degradation in prediction accuracy and the probability of assigning a sample to each class, separately. In this case, components were zeroed at the time points corresponding to the highest values in the importance vector. Specifically, for each selected group of components, the top-10 (∼ 50/512 × 100%) to top- 30 (∼ 150/512 × 100%) most important time segments were zeroed, as indicated on the X-axis of Figure 4C,D. The Y-axis represents either prediction accuracy or the probability of class assignment, respectively Classes were evaluated separately and compared with randomized zeroing of the same percentage of time segments within the same group of components, averaged over seven independently generated random samples. The results indicate that model performance for the Alzheimer’s disease class drops from 75% average accuracy at top-10 zeroed segments to 20% average accuracy at top-30 zeroed segments when zeroing the group of components corresponding to mean signal values (with 0.8 accuracy at top-0). In contrast, model performance decreases only slightly for the same class when zeroing the group of components corresponding to standard deviations, consistent with the results shown in Figure 4B. For the wild-type class, zeroing either group of components does not lead to performance degradation. Similar trends are observed in the probability of assigning an example to a given class, where the standard deviation across the entire dataset is also shown, overlaid with the standard deviation for randomized zeroing. In Supplementary Figure S5, where a comparison of different interpretability methods is also shown [31], the probability distributions at top-30 for the Alzheimer’s disease model, when zeroing the group of components with mean signal values, are statistically distinguishable from the probability distribution obtained with randomized zeroing are shown. These results suggest that the component groups corresponding to mean signal values capture the Alzheimer’s disease–related patterns present in the data, whereas the components representing standard deviations primarily provide contextual information. Consequently, the model does not simply differentiate between classes; rather, it identifies the presence of Alzheimer’s disease patterns. If such patterns are absent, the model assigns the sample to the wild-type class, consistent with the interpretation that the wild type encompasses all cases lacking disease-specific features.

## Discussion

In the present article, we introduce NEuRT, a model based on the BERT architecture, adapted to a five-component representation of calcium fluorescence intensity over time. The BERT framework was selected due to its robustness and extensive validation across various domains. Originally developed for natural language processing [29], [32], [33], it has also demonstrated strong performance in time-series analysis [34]. Building on these advantages, we propose its application for the analysis of neuronal activity patterns. Based on this rationale, we pretrained our model on the MICrONS dataset, which contains neuronal activity recordings from the mouse visual cortex, using a signal reconstruction task. The pretrained model was subsequently able to reconstruct signals from miniature fluorescence microscopy recordings obtained from the hippocampal area of both wild-type and Alzheimer’s disease model mice. Although miniature microscopy enables longitudinal recordings from freely behaving animals – a major advantage of this method – the resulting data are noisier and have lower resolution compared to two-photon microscopy. Despite these limitations, our results demonstrated that a model pretrained on a large, high-quality dataset can effectively interpret calcium imaging signals acquired with less precise techniques, across different brain regions, and even in animals exhibiting severe neurodegenerative pathology. Thus, the presented model exhibit strong generalization capabilities and hold promise for diverse neuroscience applications, particularly in scenarios where access to large datasets is limited and pretraining such models is not feasible.

It should be noted that classes have linearly separable features in the used “Miniscope” dataset. Despite this fact, NEuRT is capable of constructing a latent embedding space that accounts for multiple tasks. This suggests that in the future work, experiments could be conducted on the adapted NEuRT model for implications on the other tasks [35]. Moreover, using native NEuRT backbone as one of the encoders may find its application in a multimodal architecture for neuronal activity in combination with behavioral representations [36], [37]. It was also demonstrated that pretraining on the large dataset from the different brain region (visual cortex) obtained via another neuronal activity imaging technique (2-photon calcium imaging) improves reconstruction capabilities. Despite the linear separability between groups in the “Miniscope” dataset, which the model is prone to overfitting on [30], the interpretation performed solely on the classification task confirmed the necessity of the temporal dimension, as verified by shuffling time steps. Developed interpretation approaches for NEuRT accounts for the individual contribution of each activity matrix components. Interpretation demonstrates a key role of the components corresponding to the mean signal of the close and far neurons in distinguishing mice with the Alzheimer’s disease model. The mean value of the calcium signal reflects the overall level of neuronal activity within the population; therefore, an increase in this component can be interpreted as a higher baseline activity level, which is consistent with neuronal hyperactivity reported in Alzheimer’s disease. The standard deviation characterizes the variability of neuronal activity over time and across the local network, i.e., the spread of signal values, which may reflect differences in the stability and coordination of neuronal dynamics. It was also shown that components representing standard deviations do not have a noticeable impact on the model’s predictive capability, as demonstrated by zeroing out the top-k most important parts of the signal. However, their role becomes noticeable when complete zeroing these components causes the model to classify all samples as belonging to the Alzheimer’s disease class. This indicates that the standard deviation components provide important contextual information for classification but do not carry direct information about Alzheimer’s pathology. Considering the assumption about mean and standard deviation interpretation mentioned above, nature of the shortcuts and results from previous studies, we may assume that this effect could be correlated with neuronal hyperactivity and asynchrony correspondingly in Alzheimer’s disease rather than changes confined to individual neurons [38]–[44]. Moreover, classical statistical approach for neuronal activity quantification [3] have been validated as reliable tools for analyzing dynamic neuronal circuitry across different states [45]. Using such established analysis methods, it has been shown that 5xFAD mice exhibit increased neuronal activity, burst events, and aberrant connectivity patterns [44]. We assume that these observations are consistent with the patterns identified by the NEuRT approach in our study. The analysis of temporal segment importance was conducted using the attention rollout method. The experiment also demonstrates that zeroing out the identified time steps disrupts the temporal dependencies required by the model. At the same time, when performing top-k ablation based on the temporal segments importance (the components representing the mean signal), the model’s predictive capability for the Alzheimer’s disease class was significantly reduced. This aligns with the results of the full component ablation described above. Presumably, vital biological connections between neurons essential for coordinated hippocampus functioning are diminished after zeroing top-k temporal components leading to decreased effectiveness of the model classification. When comparing different interpretation methods, using attention pooling alone was found to provide satisfactory results, well-aligned with the findings obtained from sequential component ablation experiments. This outcome is expected as the attention pooling layer is the final stage where the linear classifier extracts class probabilities. Nevertheless, applying attention rollout further improved the localization of temporal segments associated with Alzheimer’s disease. Therefore, this method was chosen for model interpretation.

It is important to note the limitations of the presented NEuRT model and the approaches used. The main limitation of the study lies in the features nature, making the classes linearly separable. Although various approaches enable removal of the classes linear separability for logistic regression, they are leaving biologically embedded shortcuts. This issue is common in many medical studies [46]. Notably, in this work, we considered the 5xFAD model, which is characterized by pronounced alterations in neuronal activity patterns. In such conditions, the differences between classes are relatively strong and therefore easier to detect. In contrast, when addressing more subtle problems, such as distinguishing between different brain regions, behavioral states, experimental conditions, or stages of disease progression, the differences in activity are expected to be less pronounced. In these scenarios, the proposed model may be particularly useful, as it is capable of capturing more complex patterns beyond simple linear separability. Another limitation is the model’s sensitivity to the number of neurons recorded in a session. When only a small number of neurons is captured, four out of five components of the activity matrix become sparse. The approach of aggregating neuronal activity into a single activity matrix is also not ideal. While this aggregation provides flexibility regarding the number of neurons, it primarily reflects information at the macro level, and finer microscopic details may be lost. Future work may address this limitation by incorporating representations that preserve neuron-level structure or by modeling interactions between neurons more explicitly. The model also depends on hyperparameters such as learning rate and the balance of the training dataset. The presented training approach may not fully align with best practices in transformer training, but this is justified by the specifics of the problems to be solved.

In summary, the proposed NEuRT model demonstrates robustness and provides a framework for a more comprehensive and unbiased evaluation of neuronal activity patterns. In particular, it may enable a systematic and quantitative assessment of how external perturbations, including pharmacological compounds and experimental interventions, modulate neuronal activity at the level of population dynamics, by demonstrating shifts toward a healthy-like state. Notably, such an approach provides a more comprehensive and unbiased evaluation compared to analyses based on individual activity parameters, allowing for a more rigorous and interpretable assessment of intervention-induced changes in neuronal dynamics. Beyond Alzheimer’s disease models, the proposed approach may also be applicable to the study and interpretation of other neurological and pathological conditions. The methodology presented here may serve as a useful tool for developing and validating novel analytical approaches in fundamental neuroscience, particularly for the analysis and interpretation of neuronal activity patterns. This work also suggests the potential applicability of machine learning in neuroscience: while classical data analysis methods can be limited by available human resources, machine learning approaches may help to complement these techniques, potentially enabling more complex and comprehensive investigations. Looking ahead, future directions include further refinement of machine learning methods through the integration of neuronal activity with behavioral data in a multimodal framework. Another promising direction is the exploration of tasks related to the generation of neuronal activity patterns. Such approaches may enable the simulation of neuronal responses to various conditions and support in silico testing of therapeutic interventions. In addition, they may help extend the boundaries of available experimental data beyond what can be obtained in vivo.

## Supplementary Materials

All the model training experiments were conducted on two NVIDIA Tesla V100 GPUs. To ensure reproducibility, our software has been made publicly available at https://github.com/rgKric/NEuRT, along with all supplementary materials referenced in this study.

## Acknowledgment

We are thankful to all members of the Laboratory of biomedical Imaging and Data analysis and Laboratory of Molecular Neurodegeneration for the fruitful discussions. We would like to thank the head of the Laboratory of Molecular Neurodegeneration, Ilya Bezprozvanny, for providing access to valuable scientific datasets, which enables the testing of new data analysis methods.

The current study protocol was approved by the Bioethics Committee of Peter the Great St. Petersburg Polytechnic University (Ethical Permit No. 3-n-b, issued on May 25, 2022).

