## Supplemental Data 1 for "NEuRT: A Transformer-Based Model for Explainable Neuronal Activity Analysis"

### Supplementary materials

**Table S1** The number of neurons recorded in each session for WT and 5xFAD groups.

| Mouse index | session 1 | session 2 | session 3 | session 4 |
|---|---|---|---|---|
| **5xFAD** | | | | |
| 1 | 411 | 336 | — | 408 |
| 16 | 295 | 276 | — | 288 |
| 117 | 36 | 73 | 78 | 94 |
| 118 | 27 | 17 | 28 | — |
| 123 | 80 | 68 | 128 | — |
| **WT** | | | | |
| 3 | 378 | 381 | 295 | 564 |
| 5 | 413 | — | 574 | 352 |
| 7 | — | 196 | 627 | 420 |
| 102 | 173 | 72 | — | 47 |
| 106 | 106 | 341 | 82 | — |
| 109 | 144 | 97 | 125 | 108 |
| 110 | 284 | 244 | — | 275 |

#### Cross-validation

During fine-tuning, cross-validation was performed to test the hypothesis regarding the influence of individual session quality on the final results.

Cross-entropy loss of each fold is shown in Figure S1. The F1 score is shown in Figure S2. The plots are presented in a table, where each cell contains the curve for a specific fold; columns are organized by the transgenic subject ID, and rows by the wild-type subject. These plots demonstrate that the network trains stably, with the exception of some folds involving mouse 106. The average accuracy and standard deviation for each session of the validation sample calculated for the entire cross-validation experiment is shown in the Table S2.

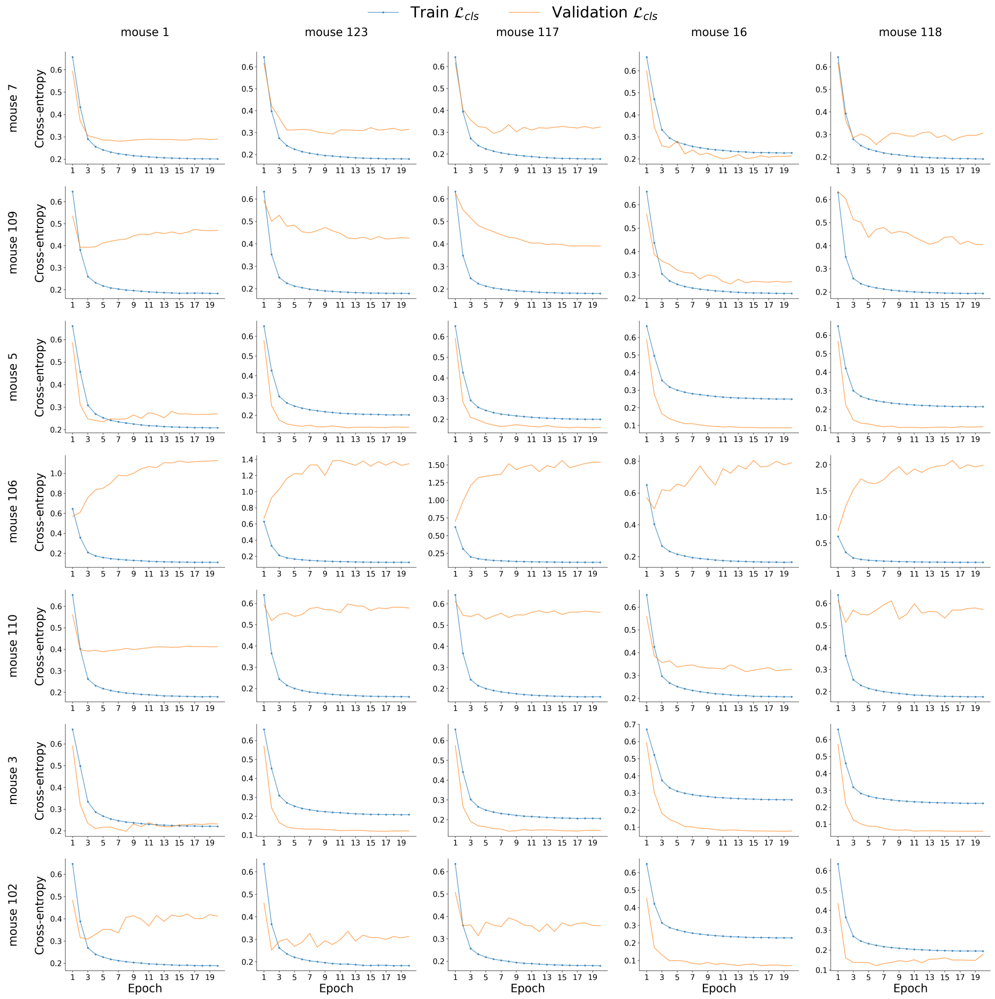


**Fig. S1** Cross-entropy loss for each fold in the cross-validation experiment. Each graph shows the loss functions by epochs for the training and validation subsample. Each row is signed on the left and corresponds to one wild-type individual. Each column is signed at the top and corresponds to a transgenic individual. The column signature and the lines of each graph show which two individuals form the same validation sample.

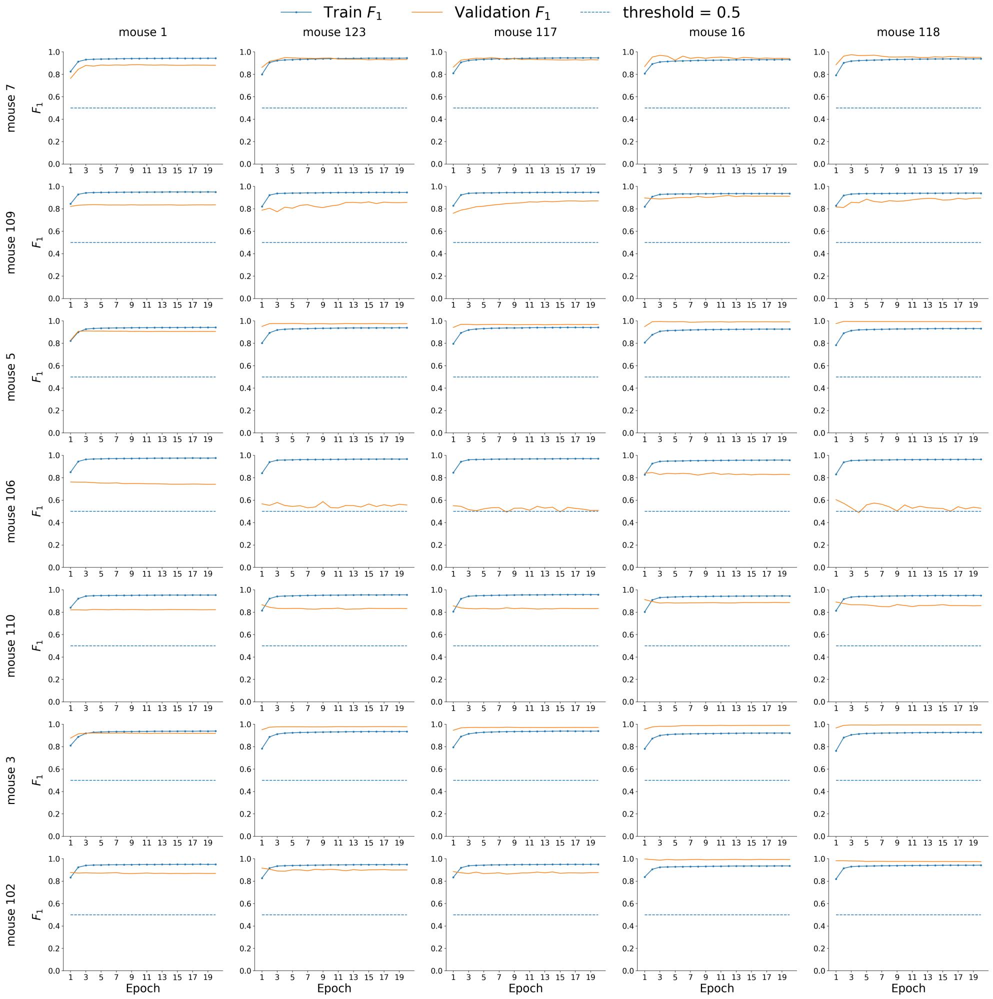


**Fig. S2** Curves of the F1 score for each fold in the cross-validation experiment. Each graph shows the F1 score by epochs for the training and validation subsample.

**Table S2** Model prediction accuracy on individual sessions for each mouse during cross-validation. Results are reported as the mean ± standard deviation across all cases in which the mouse was included in the validation set.

| Mouse index | session 1 | session 2 | session 3 | session 4 |
|---|---|---|---|---|
| **5xFAD** | | | | |
| 1 | 0.9991 ± 0.0007 | 0.2219 ± 0.0220 | — | 0.9993 ± 0.0005 |
| 16 | 0.9685 ± 0.0133 | 0.9864 ± 0.0059 | — | 1.0000 ± 0.0000 |
| 117 | 0.2682 ± 0.0869 | 0.3269 ± 0.0507 | 0.9905 ± 0.0046 | 0.9174 ± 0.0426 |
| 118 | 0.6092 ± 0.0674 | 0.3352 ± 0.1178 | 0.9982 ± 0.0025 | — |
| 123 | 0.7880 ± 0.0299 | 0.2997 ± 0.0789 | 0.9993 ± 0.0007 | — |
| **WT** | | | | |
| 3 | 1.0000 ± 0.0000 | 1.0000 ± 0.0000 | 0.9901 ± 0.0096 | 0.9973 ± 0.0008 |
| 5 | 0.9980 ± 0.0020 | — | 0.9971 ± 0.0021 | 1.0000 ± 0.0000 |
| 7 | — | 0.6449 ± 0.0493 | 0.9799 ± 0.0034 | 0.9433 ± 0.0136 |
| 102 | 1.0000 ± 0.0000 | 1.0000 ± 0.0000 | — | 1.0000 ± 0.0000 |
| 106 | 0.8282 ± 0.0625 | 0.2150 ± 0.0320 | 0.4926 ± 0.0448 | — |
| 109 | 0.8042 ± 0.0795 | 0.8580 ± 0.0597 | 0.8864 ± 0.0708 | 0.6930 ± 0.0811 |
| 110 | 0.9986 ± 0.0015 | 0.9989 ± 0.0017 | — | 0.3175 ± 0.0328 |

In this experiment, 0.5 was chosen as the threshold value for the following reasons. Each activity matrix actually represents the activity of one neuron, provided with some additional statistics about its environment. Thus, the proportion of matrices obtained from one session to a specific class can act as an indirect metric of the model's confidence in predicting the entire session. If more than half of the data from one session is assigned by the model to a specific class, then it is classified to this class.

For all folds used in cross-validation, we plotted the evolution of prediction accuracy for each individual mouse session in the validation set. The resulting graph is shown in Figure S3.

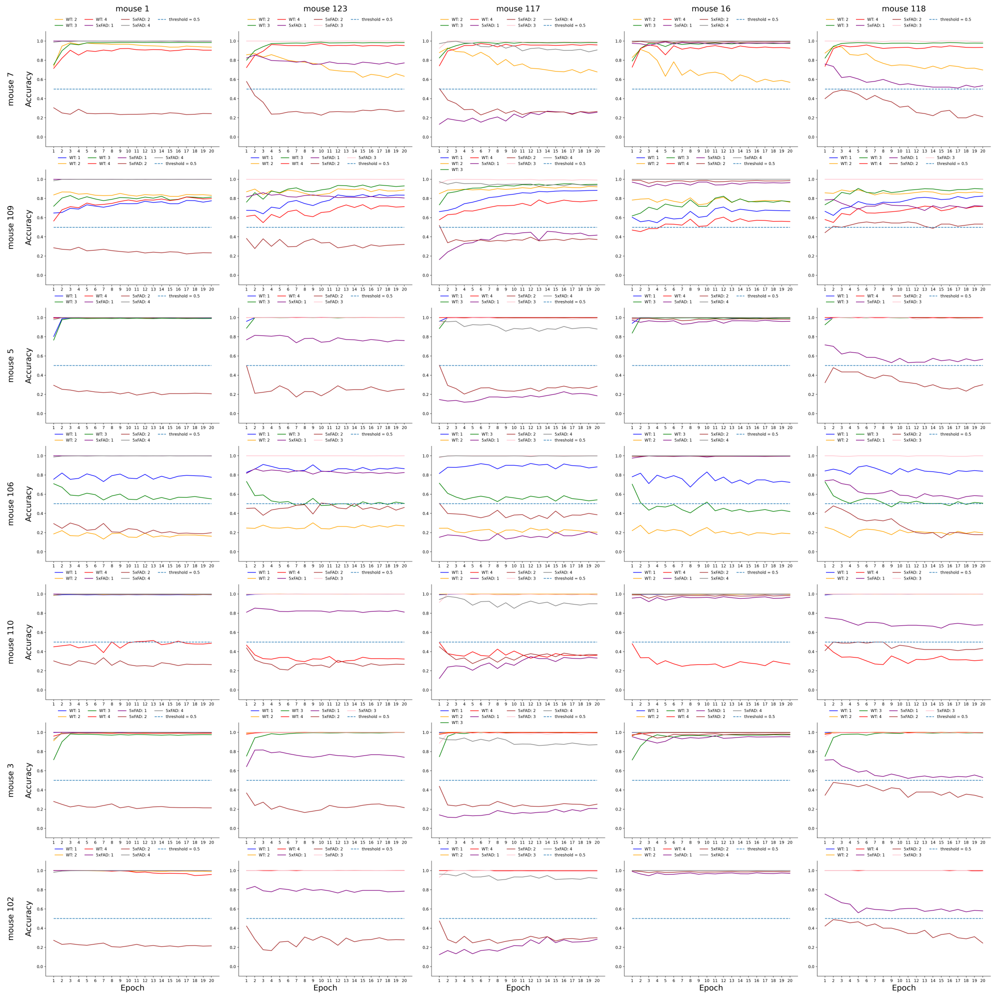


**Fig. S3** Classification accuracy across validation sessions for each fold. Each graph shows the dependence of classification accuracy on the epoch number. The threshold is equal to 0.5.

In the cross-validation experiment, a multitask learning approach was employed. Table S3 presents the final metrics obtained from the cross-validation experiment for the validation set.

**Table S3** Cross-validation performance metrics for the validation set using a multitask learning approach. The table shows the results for each combination of mouse indices from the wild-type (WT) and 5xFAD groups.

| WT | 5xFAD | $R^2$ | MAE | cosine similarity | precision | recall |
|---|---|---|---|---|---|---|
| 7 | 1 | $0.9868 \pm 0.0049$ | $0.0051 \pm 0.0024$ | $0.9922 \pm 0.0140$ | 0.9499 | 0.8224 |
| 7 | 123 | $0.9794 \pm 0.0056$ | $0.0052 \pm 0.0024$ | $0.9927 \pm 0.0121$ | 0.9182 | 0.9464 |
| 7 | 117 | $0.9783 \pm 0.0054$ | $0.0051 \pm 0.0023$ | $0.9922 \pm 0.0131$ | 0.9269 | 0.9316 |
| 7 | 16 | $0.9852 \pm 0.0053$ | $0.0052 \pm 0.0026$ | $0.9930 \pm 0.0123$ | 0.8949 | 0.9929 |
| 7 | 118 | $0.9786 \pm 0.0053$ | $0.0050 \pm 0.0023$ | $0.9923 \pm 0.0129$ | 0.9190 | 0.9858 |
| 109 | 1 | $0.9821 \pm 0.0047$ | $0.0061 \pm 0.0025$ | $0.9921 \pm 0.0125$ | 0.8028 | 0.5944 |
| 109 | 123 | $0.9647 \pm 0.0085$ | $0.0076 \pm 0.0024$ | $0.9927 \pm 0.0080$ | 0.8437 | 0.8713 |
| 109 | 117 | $0.9613 \pm 0.0102$ | $0.0075 \pm 0.0023$ | $0.9919 \pm 0.0096$ | 0.8849 | 0.8578 |
| 109 | 16 | $0.9782 \pm 0.0049$ | $0.0064 \pm 0.0027$ | $0.9932 \pm 0.0098$ | 0.6900 | 0.9579 |
| 109 | 118 | $0.9553 \pm 0.0128$ | $0.0081 \pm 0.0023$ | $0.9920 \pm 0.0076$ | 0.8272 | 0.9755 |
| 5 | 1 | $0.9783 \pm 0.0080$ | $0.0060 \pm 0.0026$ | $0.9923 \pm 0.0126$ | 0.9943 | 0.8322 |
| 5 | 123 | $0.9576 \pm 0.0132$ | $0.0066 \pm 0.0026$ | $0.9927 \pm 0.0103$ | 0.9991 | 0.9525 |
| 5 | 117 | $0.9533 \pm 0.0143$ | $0.0065 \pm 0.0025$ | $0.9923 \pm 0.0111$ | 0.9993 | 0.9380 |
| 5 | 16 | $0.9739 \pm 0.0095$ | $0.0061 \pm 0.0027$ | $0.9931 \pm 0.0108$ | 0.9957 | 0.9878 |
| 5 | 118 | $0.9474 \pm 0.0158$ | $0.0066 \pm 0.0026$ | $0.9924 \pm 0.0107$ | 0.9986 | 0.9888 |
| 106 | 1 | $0.9851 \pm 0.0048$ | $0.0058 \pm 0.0026$ | $0.9914 \pm 0.0152$ | 0.3412 | 0.3965 |
| 106 | 123 | $0.9751 \pm 0.0073$ | $0.0062 \pm 0.0027$ | $0.9923 \pm 0.0119$ | 0.4217 | 0.8201 |
| 106 | 117 | $0.9730 \pm 0.0080$ | $0.0061 \pm 0.0026$ | $0.9913 \pm 0.0140$ | 0.3881 | 0.7422 |
| 106 | 16 | $0.9825 \pm 0.0054$ | $0.0058 \pm 0.0029$ | $0.9927 \pm 0.0125$ | 0.3283 | 0.9901 |
| 106 | 118 | $0.9735 \pm 0.0079$ | $0.0061 \pm 0.0027$ | $0.9914 \pm 0.0134$ | 0.3696 | 0.9240 |
| 110 | 1 | $0.9838 \pm 0.0057$ | $0.0057 \pm 0.0025$ | $0.9924 \pm 0.0129$ | 0.8202 | 0.7261 |
| 110 | 123 | $0.9714 \pm 0.0095$ | $0.0064 \pm 0.0024$ | $0.9933 \pm 0.0096$ | 0.7677 | 0.9086 |
| 110 | 117 | $0.9687 \pm 0.0102$ | $0.0063 \pm 0.0023$ | $0.9928 \pm 0.0106$ | 0.7815 | 0.8917 |
| 110 | 16 | $0.9808 \pm 0.0068$ | $0.0059 \pm 0.0027$ | $0.9936 \pm 0.0103$ | 0.7473 | 0.9771 |
| 110 | 118 | $0.9670 \pm 0.0110$ | $0.0064 \pm 0.0023$ | $0.9932 \pm 0.0095$ | 0.7640 | 0.9815 |
| 3 | 1 | $0.9853 \pm 0.0059$ | $0.0054 \pm 0.0022$ | $0.9892 \pm 0.0187$ | 0.9913 | 0.8574 |
| 3 | 123 | $0.9764 \pm 0.0075$ | $0.0057 \pm 0.0021$ | $0.9885 \pm 0.0186$ | 0.9988 | 0.9579 |
| 3 | 117 | $0.9748 \pm 0.0075$ | $0.0056 \pm 0.0020$ | $0.9881 \pm 0.0195$ | 0.9984 | 0.9467 |
| 3 | 16 | $0.9833 \pm 0.0063$ | $0.0055 \pm 0.0023$ | $0.9896 \pm 0.0175$ | 0.9939 | 0.9881 |
| 3 | 118 | $0.9737 \pm 0.0074$ | $0.0057 \pm 0.0020$ | $0.9871 \pm 0.0210$ | 0.9979 | 0.9905 |
| 102 | 1 | $0.9880 \pm 0.0047$ | $0.0055 \pm 0.0023$ | $0.9882 \pm 0.0203$ | 0.9905 | 0.5167 |
| 102 | 123 | $0.9779 \pm 0.0100$ | $0.0065 \pm 0.0021$ | $0.9829 \pm 0.0263$ | 1.0000 | 0.8185 |
| 102 | 117 | $0.9745 \pm 0.0116$ | $0.0062 \pm 0.0019$ | $0.9819 \pm 0.0283$ | 1.0000 | 0.7800 |
| 102 | 16 | $0.9857 \pm 0.0060$ | $0.0057 \pm 0.0026$ | $0.9892 \pm 0.0177$ | 1.0000 | 0.9596 |
| 102 | 118 | $0.9678 \pm 0.0155$ | $0.0065 \pm 0.0019$ | $0.9757 \pm 0.0373$ | 1.0000 | 0.9491 |

### Baseline Experiment

To evaluate the performance of the presented model, we conducted a comparison against baseline: a recurrent neural network (RNN), a logistic regression classifier. Also, the final version of NEuRT performance was compared with itself without pre-train stage. The RNN baseline shared the same output head and dimensionality as our main model. The logistic regression baseline was implemented by flattening the input channels ($512 \times 5$) into a single vector of size 2560, followed by a linear layer without activation.

**Table S4** Models' performance comparison on the reconstruction and classification tasks. The same test sample and seed was used for all models test. The last training epoch results were used.

| | Reconstruction | | | | |
|---|---|---|---|---|---|
| Channel | Signal | $\mu_{\text{close}}$ | $\sigma_{\text{close}}$ | $\mu_{\text{far}}$ | $\sigma_{\text{far}}$ |
| | | | $\mathbf{R^2}$ | | |
| NEuRT with pretrain | 0.9766 | 0.9827 | 0.9595 | 0.9836 | 0.9662 |
| NEuRT without pretrain | 0.4918 | 0.6398 | 0.3901 | 0.7284 | 0.4120 |
| RNN | 0.7535 | 0.0015 | -0.0650 | -0.0434 | -0.0729 |
| LogisticRegression | NaN | NaN | NaN | NaN | NaN |
| | | | **MAE** | | |
| NEuRT with pretrain | 0.0112 | 0.0047 | 0.0065 | 0.0036 | 0.0046 |
| NEuRT without pretrain | 0.0893 | 0.0277 | 0.0334 | 0.0185 | 0.0257 |
| RNN | 0.0607 | 0.0471 | 0.0433 | 0.0365 | 0.0330 |
| LogisticRegression | NaN | NaN | NaN | NaN | NaN |
| | | | **cosine similarity** | | |
| NEuRT with pretrain | 0.9714 | 0.9982 | 0.9976 | 0.9990 | 0.9990 |
| NEuRT without pretrain | 0.6589 | 0.9646 | 0.9664 | 0.9865 | 0.9848 |
| RNN | 0.8089 | 0.9675 | 0.9689 | 0.9881 | 0.9863 |
| LogisticRegression | NaN | NaN | NaN | NaN | NaN |
| | | | Classification | | |
| | | **precision** | | **recall** | |
| NEuRT with pretrain | | 0.9948 | | 0.9850 | |
| NEuRT without pretrain | | 0.7875 | | 0.8790 | |
| RNN | | 0.6450 | | 0.6488 | |
| LogisticRegression | | 0.9963 | | 0.9507 | |

The baseline experiment highlights the comparative performance of the RNN, logistic regression, and NEuRT models across reconstruction and classification tasks. Table S4 demonstrates that the NEuRT model with pretraining consistently outperforms the other baselines, achieving the highest scores in $R^2$, MAE, and cosine similarity.

#### Interpretation methods

In Figure S4, the confusion matrices obtained by zeroing out individual components of the activity matrix are shown.

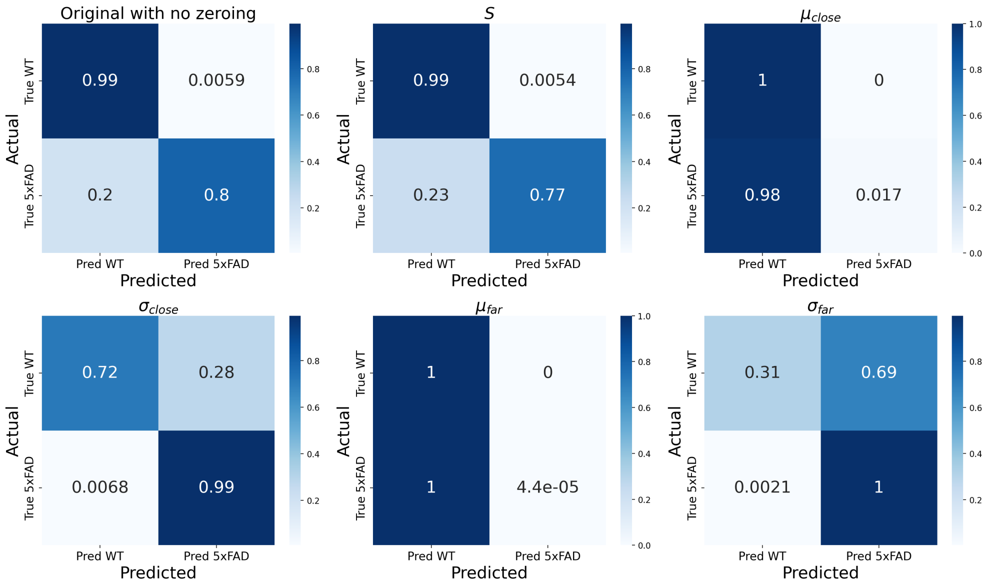


**Fig. S4** Confusion matrices on the slice level for the model outputs with zeroing activity matrix's individual components. The color gradient indicates results normalized to the true value.

To estimate the importance of temporal segments, we employed an adapted attention rollout method, while also testing simplified and extended variants. Two modifications were considered. The first involved using entropy within a single attention layer to reduce the dominance of diagonal attention matrices:

$$W_l = \text{softmax}(\frac{1}{L}\sum_{i=1}^{L}(-\sum_{j=1}^{L} A_{lij}^h \log_2 (A_{lij}^h + 10^{-12}))$$
$$\overline{A_l^h} = W_l^h \cdot A_l^h$$

Instead of the standard averaging over attention heads within a layer, a weighted average was applied.

The second modification involved using gradients as weights, not for entire matrices, but for individual elements. A similar approach was used in (Chefer et al., 2021), where relevance weights were applied instead of attention matrices. The gradient-based weighting is expressed as:

$$\overline{A_l^h} = \nabla A_l^h \odot A_l^h$$

We compared variants using entropy alone and entropy combined with gradients. The most notable behavior, however, was observed when using only the pooling vector weights to estimate temporal importance, as shown in Figure S5, where methods are compared in the top-k zeroing experiments described in the Results.

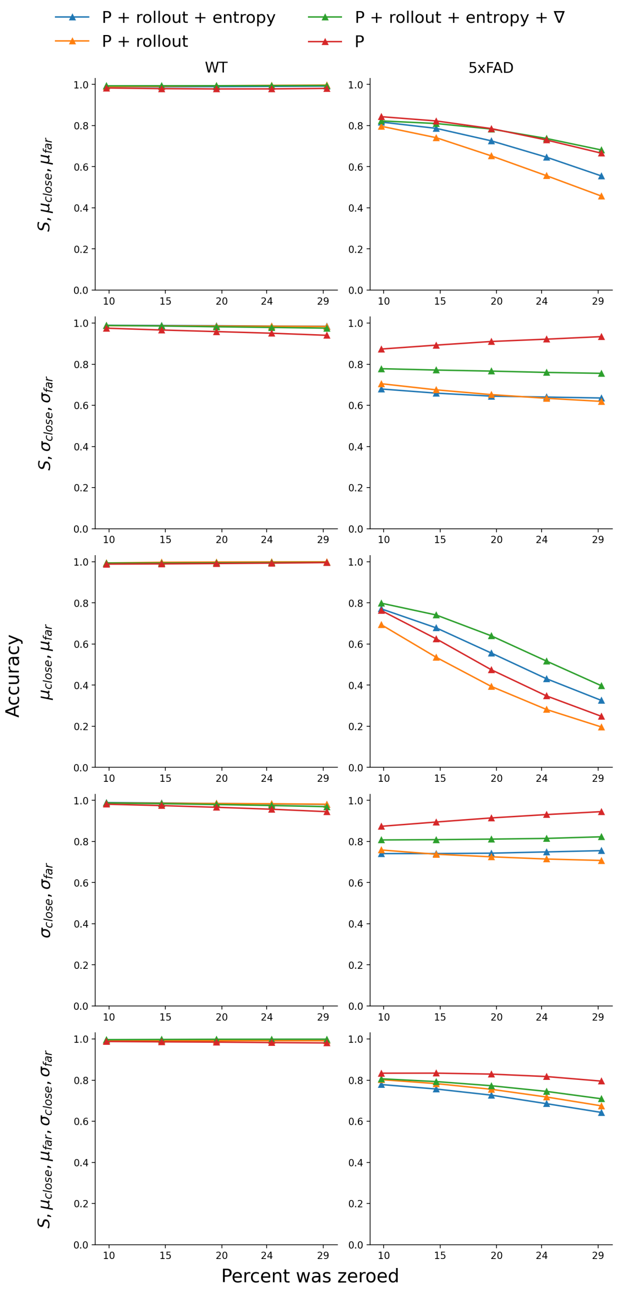


**Fig. S5** Comparison of the model's prediction accuracy for both classes when zeroing out the top-k temporal segments of activity matrix component groups based on their importance using different interpretation methods: P + rollout + entropy – a combination of pooling weights and entropy-weighted attention rollout; P + rollout – a combination of pooling weights and unweighted attention rollout; P + rollout + entropy + ∇ – a combination of pooling weights, entropy-weighted attention rollout, and gradients; P – pooling weights alone.
